## Supplementary material for "Auto QSAR-based Active learning docking for hit identification of potential inhibitors of *Plasmodium falciparum* Hsp90 as antimalarial agents": Supplimentary data: 17May2024_Final_Supplementary data_AL paper (LM).docx

A. **Reaction based enumeration**

**A**

**B**

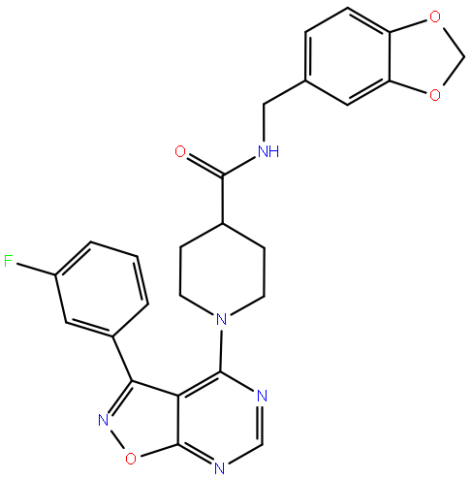

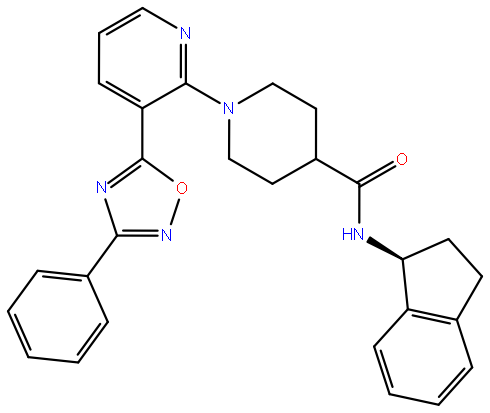

**Figure S1**: reference compounds from the Everson et al 2021 study. A being compound 7 and B being compound 10

**Table S1**:Pathfinder generated reaction pathways -Schrödinger Suite 2019-1

| Path 1 | Amide_coupling-1 |
| --- | --- |
| Path 2 | Amide_coupling-2 |
| Path 3 | Amination-1 |
| Path 4 | Hiyama-1 |
| Path 5 | Negishi |
| Path 6 | Oxadiazole-1 |
| Path 7 | Stille |
| Path 8 | Suzuki |
| Path 9 | Suzuki-2 |

Pathway 6 which is oxadiazole-1 was chosen our pathway of interest shown in figure 1 below

### B. **AutoQSAR model generation**

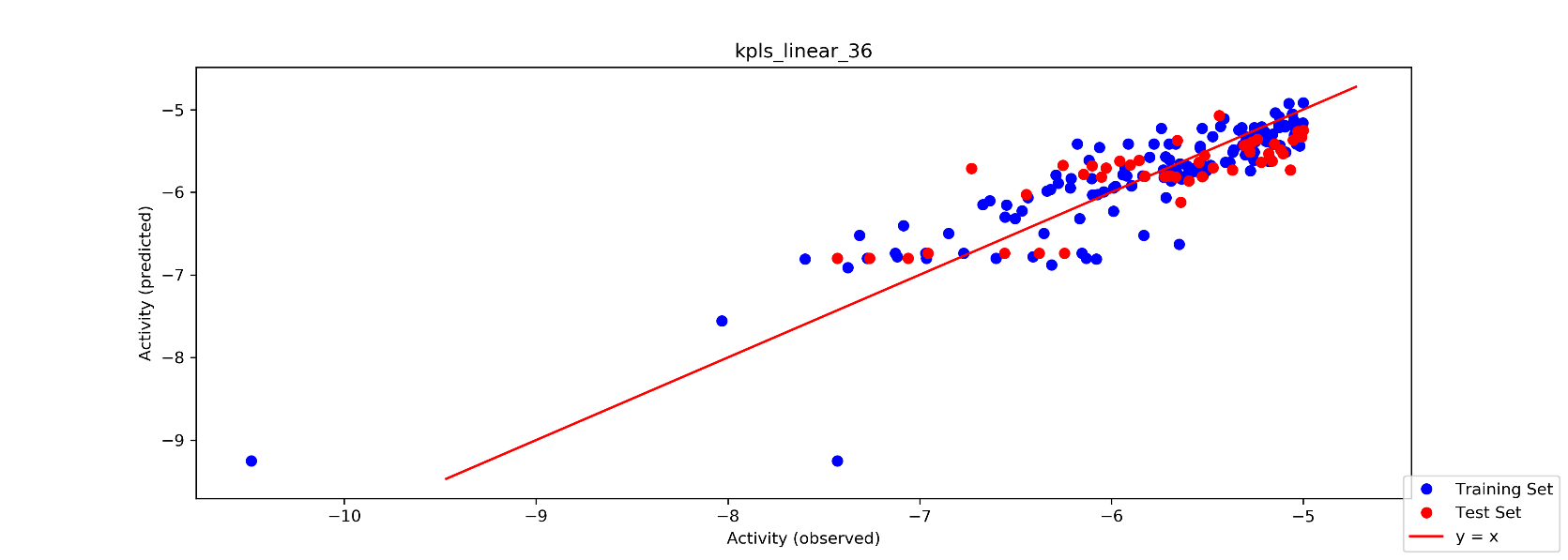

**Figure S2**: model 1

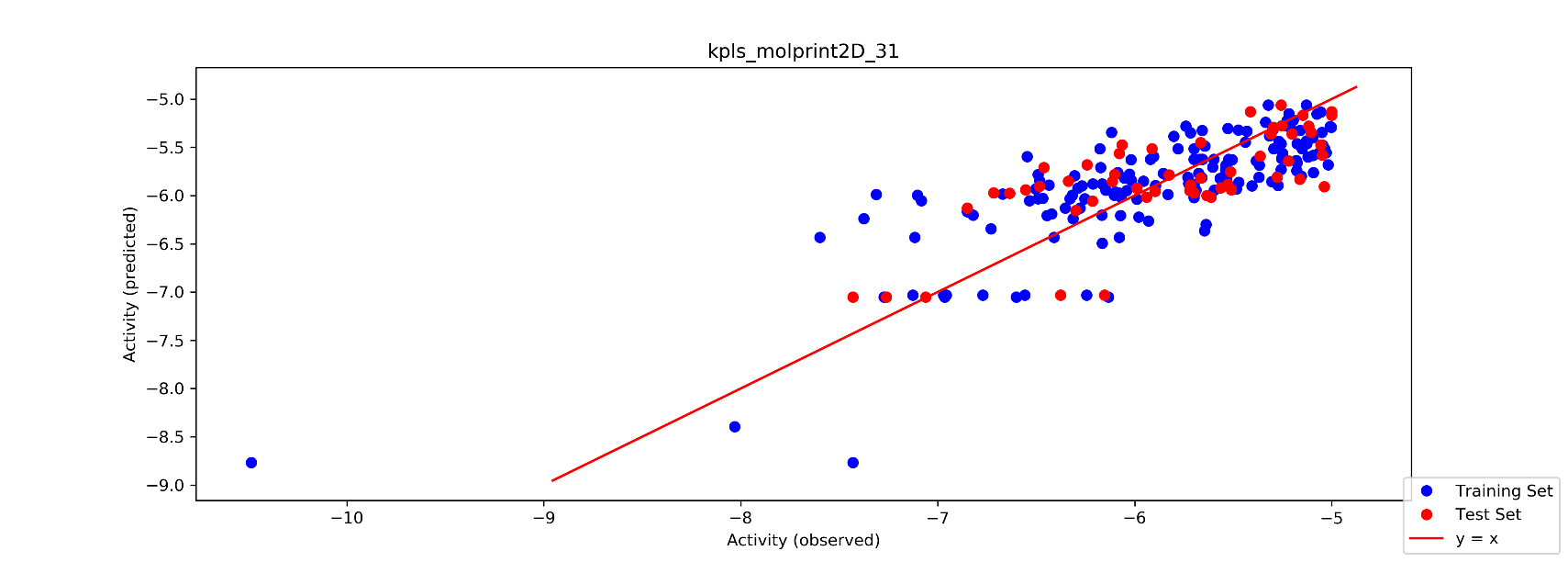

**Figure S3**:model2

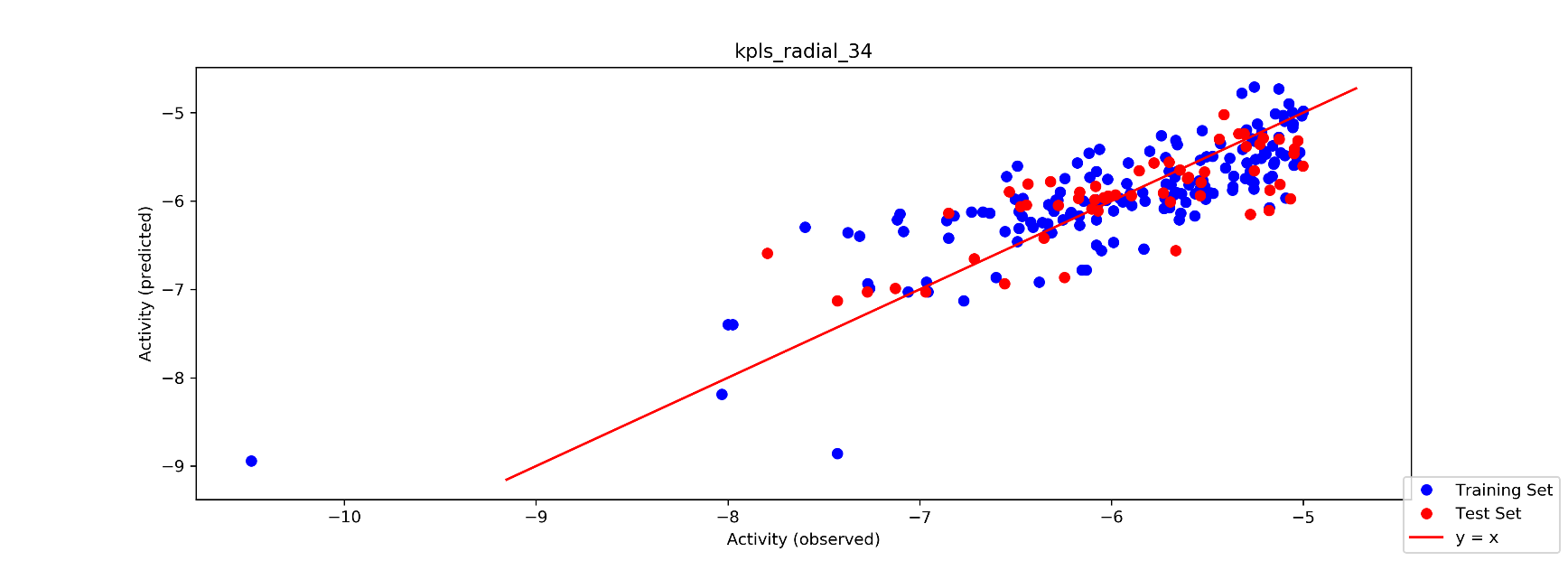

**Figure S4**: model 3

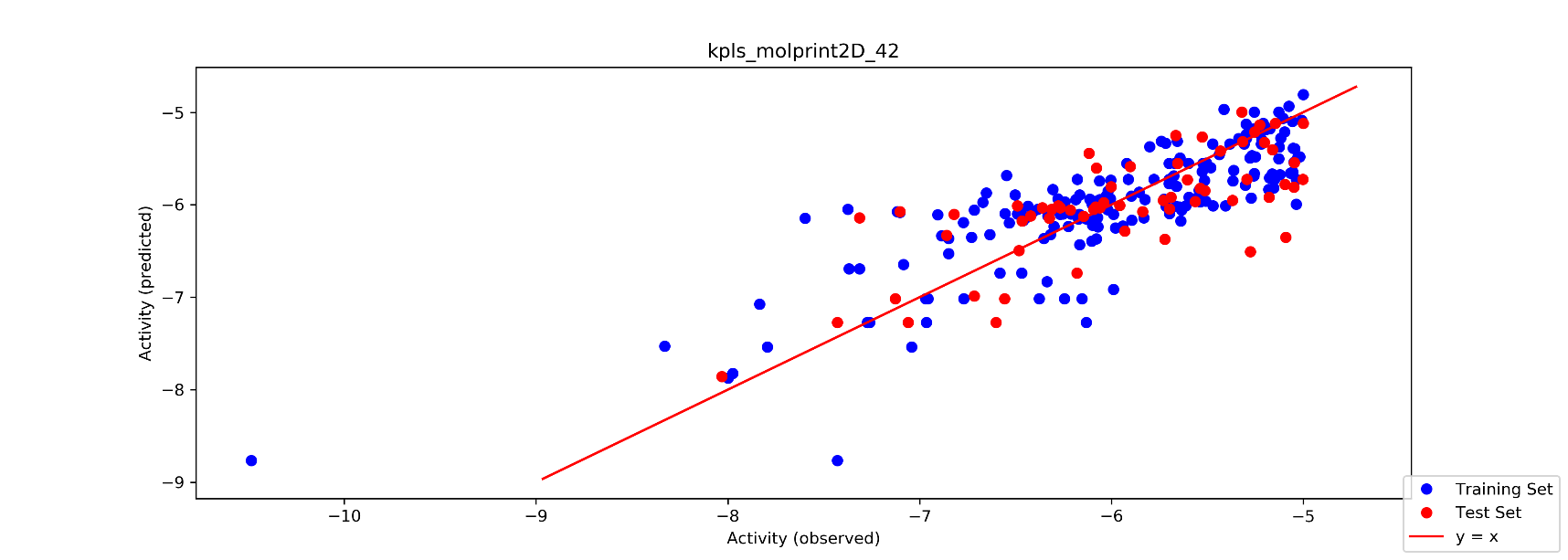

**Figure S5**:model4

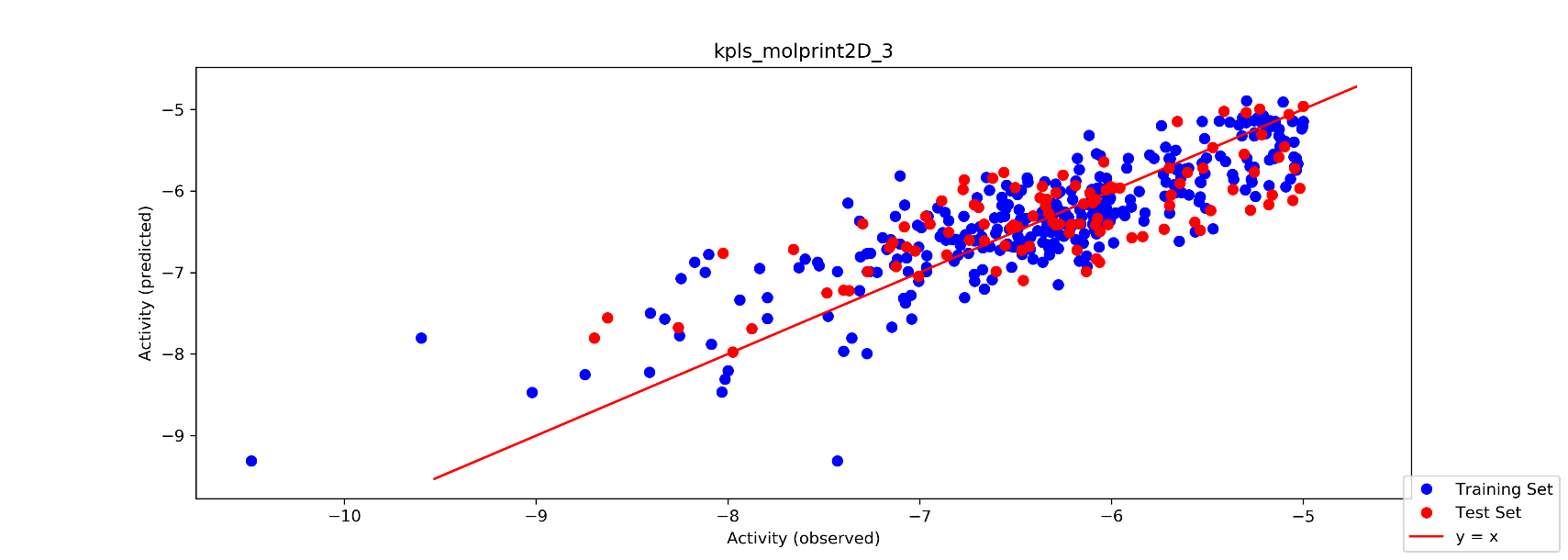

**Figure S6**: model 10

### C. **Induced fit docking**

**Table S2:** induced fit docking results of the top 62 compounds against the PfHsp90 N-terminal domain receptor

| Title | docking score | glide gscore | glide emodel | XP GScore | IFDScore |
| --- | --- | --- | --- | --- | --- |
| FTN40281232 | -12.032 | -12.058 | -106.600 | -12.058 | -526.97 |
| FTN 2 | -12.351 | -12.351 | -126.680 | -12.351 | -526.86 |
| FTN315699906 | -10.501 | -10.746 | -113.012 | -10.746 | -526.14 |
| FTN315452139 | -10.396 | -10.396 | -117.889 | -10.396 | -525.99 |
| FTN48606212 | -10.796 | -10.796 | -92.753 | -10.796 | -524.35 |
| FTN 3 | -10.380 | -10.457 | -101.975 | -10.457 | -523.50 |
| FTN5 | -11.216 | -11.216 | -103.704 | -11.216 | -523.34 |
| FTN39249872 | -10.178 | -10.431 | -65.285 | -10.431 | -523.26 |
| FTN41168581 | -11.209 | -11.224 | -84.635 | -11.224 | -522.53 |
| FTN313761635 | -10.603 | -10.611 | -86.084 | -10.611 | -520.29 |
| FTN4919 | -13.161 | -13.162 | -108.122 | -13.162 | -533.02 |
| FTN717 | -12.399 | -12.435 | -85.494 | -12.435 | -528.05 |
| FTN1424 | -11.975 | -11.976 | -107.794 | -11.976 | -526.61 |
| FTN1841 | -11.396 | -11.885 | -105.884 | -11.885 | -526.20 |
| FTN1424 | -12.446 | -12.447 | -101.188 | -12.447 | -526.03 |
| FTN 1 | -14.746 | -14.794 | -120.018 | -14.794 | -525.66 |
| FTN3967 | -11.880 | -12.442 | -111.528 | -12.442 | -525.22 |
| FTN4079 | -13.233 | -14.746 | -116.205 | -14.746 | -524.84 |
| FTN1133 | -11.006 | -11.171 | -107.716 | -11.171 | -524.79 |
| FTN 4 | -14.313 | -14.313 | -105.919 | -14.313 | -524.73 |
| FTN1133 | -10.175 | -11.013 | -106.508 | -11.013 | -524.43 |
| FTN1841 | -11.894 | -12.235 | -86.219 | -12.235 | -524.04 |
| FTN1715 | -10.790 | -13.133 | -102.921 | -13.133 | -523.90 |
| FTN3630 | -13.914 | -13.914 | -84.466 | -13.914 | -523.58 |
| FTN1813 | -11.941 | -12.423 | -101.299 | -12.423 | -523.44 |
| FTN4994 | -10.301 | -10.301 | -101.453 | -10.301 | -523.43 |
| FTN3885 | -13.873 | -13.874 | -100.826 | -13.874 | -522.27 |
| FTN4184 | -10.790 | -10.955 | -102.957 | -10.955 | -522.18 |
| FTN4363 | -12.400 | -12.400 | -90.507 | -12.400 | -522.02 |
| FTN3782 | -10.461 | -10.461 | -103.342 | -10.461 | -522.00 |
| FTN1261 | -12.330 | -12.332 | -98.280 | -12.332 | -521.62 |
| FTN3885 | -12.396 | -12.397 | -105.667 | -12.397 | -521.51 |
| FTN499 | -12.397 | -12.759 | -89.158 | -12.759 | -521.49 |
| FTN4363 | -12.281 | -12.281 | -92.724 | -12.281 | -521.47 |
| FTN1715 | -13.770 | -13.781 | -112.874 | -13.781 | -521.33 |
| FTN4843 | -11.544 | -11.544 | -85.764 | -11.544 | -521.27 |
| FTN499 | -11.470 | -11.933 | -115.374 | -11.933 | -521.21 |
| FTN3885 | -11.725 | -11.726 | -105.179 | -11.726 | -521.15 |
| FTN3885 | -11.995 | -11.996 | -108.696 | -11.996 | -521.07 |
| FTN2075 | -11.412 | -11.983 | -117.573 | -11.983 | -520.93 |
| FTN2925 | -10.489 | -11.022 | -119.488 | -11.022 | -520.92 |
| FTN3057 | -11.430 | -11.430 | -78.680 | -11.430 | -520.64 |
| FTN2701 | -10.676 | -10.774 | -104.290 | -10.774 | -520.06 |
| FTN4031 | -10.535 | -10.535 | -91.366 | -10.535 | -519.80 |
| FTN2474 | -12.242 | -12.242 | -89.123 | -12.242 | -519.66 |
| FTN0 | -11.056 | -11.056 | -77.271 | -11.056 | -519.33 |
| FTN2110 | -12.134 | -12.153 | -76.051 | -12.153 | -519.33 |
| FTN1589 | -10.151 | -10.151 | -79.481 | -10.151 | -518.91 |
| FTN2211 | -10.254 | -10.264 | -79.071 | -10.264 | -518.87 |
| FTN1388 | -10.186 | -10.188 | -73.541 | -10.188 | -517.80 |
| FTN1454 | -10.579 | -11.025 | -80.285 | -11.025 | -517.77 |
| FTN3550 | -11.025 | -11.025 | -52.640 | -11.025 | -517.74 |
| FTN3550 | -11.156 | -11.156 | -75.256 | -11.156 | -517.20 |
| FTN3967 | -11.988 | -12.278 | -89.206 | -12.278 | -516.97 |
| FTN3885 | -14.593 | -14.594 | -96.499 | -14.594 | -516.89 |
| FTN3550 | -10.190 | -10.190 | -54.918 | -10.190 | -516.65 |
| FTN1956 | -12.433 | -12.433 | -56.229 | -12.433 | -516.10 |
| FTN4541 | -10.567 | -10.567 | -82.188 | -10.567 | -515.48 |
| FTN2403 | -14.162 | -14.162 | -100.693 | -14.162 | -515.14 |
| FTN3816 | -12.238 | -12.238 | -71.732 | -12.238 | -514.82 |
| FTN95 | -12.502 | -12.502 | -88.717 | -12.502 | -514.19 |
| FTN 6 | -14.176 | -14.176 | -77.866 | -14.176 | -512.26 |

**Human Hsp90 IFD results**

**Table S3** IFD Results of top 62 docked ligands against human Hsp90

| Title | docking score | glide gscore | glide emodel | XP GScore | IFDScore |
| --- | --- | --- | --- | --- | --- |
| FTN 315452139 | -12.108 | -12.108 | -107.792 | -12.108 | -483.60 |
| FTN 4919 | -8.588 | -8.589 | -74.147 | -8.589 | -483.16 |
| FTN 717 | -11.344 | -11.381 | -69.257 | -11.381 | -481.77 |
| FTN 4843 | -8.858 | -8.858 | -69.833 | -8.858 | -480.61 |
| FTN 4994 | -11.274 | -11.274 | -94.124 | -11.274 | -481.07 |
| FTN 40281232 | -10.106 | -10.132 | -119.206 | -10.132 | -480.67 |
| FTN 3967 | -9.843 | -10.405 | -88.841 | -10.405 | -480.67 |
| FTN 1841 | -10.009 | -10.498 | -81.878 | -10.498 | -480.48 |
| FTN 1424 | -7.406 | -7.407 | -88.461 | -7.407 | -480.47 |
| FTN 1424 | -8.161 | -8.162 | -73.416 | -8.162 | -480.20 |
| FTN 3967 | -10.750 | -11.040 | -70.525 | -11.040 | -479.85 |
| FTN 499 | -11.241 | -11.705 | -73.851 | -11.705 | -478.94 |
| FTN 1 | -9.696 | -11.208 | -83.033 | -11.208 | -479.05 |
| FTN 3782 | -9.887 | -9.887 | -93.335 | -9.887 | -479.01 |
| FTN 4079 | -10.721 | -10.769 | -84.547 | -10.769 | -478.69 |
| FTN 39249872 | -9.112 | -9.365 | -101.781 | -9.365 | -478.89 |
| FTN 315699906 | -8.438 | -8.683 | -110.313 | -8.683 | -478.87 |
| FTN 4184 | -9.628 | -9.793 | -78.937 | -9.793 | -478.49 |
| FTN 48606212 | -8.840 | -8.840 | -96.472 | -8.840 | -478.70 |
| FTN 1813 | -9.635 | -10.117 | -86.519 | -10.117 | -478.67 |
| FTN 1133 | -9.546 | -9.711 | -76.544 | -9.711 | -477.99 |
| FTN 1133 | -7.736 | -8.573 | -88.841 | -8.573 | -477.97 |
| FTN 3885 | -11.935 | -11.936 | -88.061 | -11.936 | -477.97 |
| FTN 3885 | -12.258 | -12.259 | -90.306 | -12.259 | -477.85 |
| FTN 4363 | -10.540 | -10.540 | -70.817 | -10.540 | -477.73 |
| FTN 1841 | -9.820 | -10.162 | -61.125 | -10.162 | -477.44 |
| FTN 499 | -9.933 | -10.295 | -69.552 | -10.295 | -477.60 |
| FTN 1715 | -10.042 | -10.054 | -80.589 | -10.054 | -477.40 |
| FTN 1715 | -7.067 | -9.410 | -64.766 | -9.410 | -476.95 |
| FTN 2 | -7.718 | -7.718 | -91.858 | -7.718 | -477.49 |
| FTN 3 | -8.091 | -8.168 | -69.566 | -8.168 | -476.92 |
| FTN 1454 | -7.794 | -8.240 | -60.846 | -8.240 | -477.18 |
| FTN 3885 | -11.247 | -11.248 | -81.836 | -11.248 | -477.02 |
| FTN 313761635 | -9.566 | -9.574 | -91.269 | -9.574 | -476.93 |
| FTN 4 | -10.573 | -10.573 | -80.879 | -10.573 | -476.88 |
| FTN 0 | -10.606 | -10.606 | -73.063 | -10.606 | -476.82 |
| FTN 41168581 | -9.639 | -9.653 | -97.019 | -9.653 | -476.80 |
| FTN 5 | -9.291 | -9.291 | -100.868 | -9.291 | -476.79 |
| FTN 3885 | -11.205 | -11.206 | -85.156 | -11.206 | -476.64 |
| FTN 3885 | -11.356 | -11.357 | -82.290 | -11.357 | -476.22 |
| FTN 4363 | -9.320 | -9.320 | -71.521 | -9.320 | -476.12 |
| FTN 2701 | -9.197 | -9.295 | -58.078 | -9.295 | -475.78 |
| FTN 2075 | -8.427 | -8.997 | -88.691 | -8.997 | -475.72 |
| FTN 2925 | -7.917 | -8.450 | -74.441 | -8.450 | -474.97 |
| FTN 4031 | -8.559 | -8.559 | -67.429 | -8.559 | -475.17 |
| FTN 4541 | -7.923 | -7.923 | -64.967 | -7.923 | -475.04 |
| FTN 3630 | -9.246 | -9.246 | -66.496 | -9.246 | -474.60 |
| FTN 6 | -9.476 | -9.476 | -64.259 | -9.476 | -474.34 |
| FTN 1956 | -8.851 | -8.851 | -70.780 | -8.851 | -474.27 |
| FTN 2474 | -8.353 | -8.353 | -63.745 | -8.353 | -474.23 |
| FTN 1261 | -8.915 | -8.917 | -80.463 | -8.917 | -474.18 |
| FTN 3057 | -8.137 | -8.137 | -54.889 | -8.137 | -474.07 |
| FTN 2110 | -8.201 | -8.220 | -62.872 | -8.220 | -473.94 |
| FTN 2211 | -8.205 | -8.215 | -50.809 | -8.215 | -472.91 |
| FTN 1589 | -7.292 | -7.292 | -57.357 | -7.292 | -473.34 |
| FTN 1388 | -7.042 | -7.044 | -43.262 | -7.044 | -473.23 |
| FTN 3550 | -9.021 | -9.021 | -64.583 | -9.021 | -473.18 |
| FTN 2403 | -8.844 | -8.844 | -61.754 | -8.844 | -472.83 |
| FTN 3816 | -8.956 | -8.956 | -54.034 | -8.956 | -472.53 |
| FTN 3550 | -8.071 | -8.071 | -57.252 | -8.071 | -472.15 |
| FTN 3550 | -8.064 | -8.064 | -54.193 | -8.064 | -471.96 |
| FTN 95 | -8.160 | -8.160 | -57.789 | -8.160 | -470.98 |

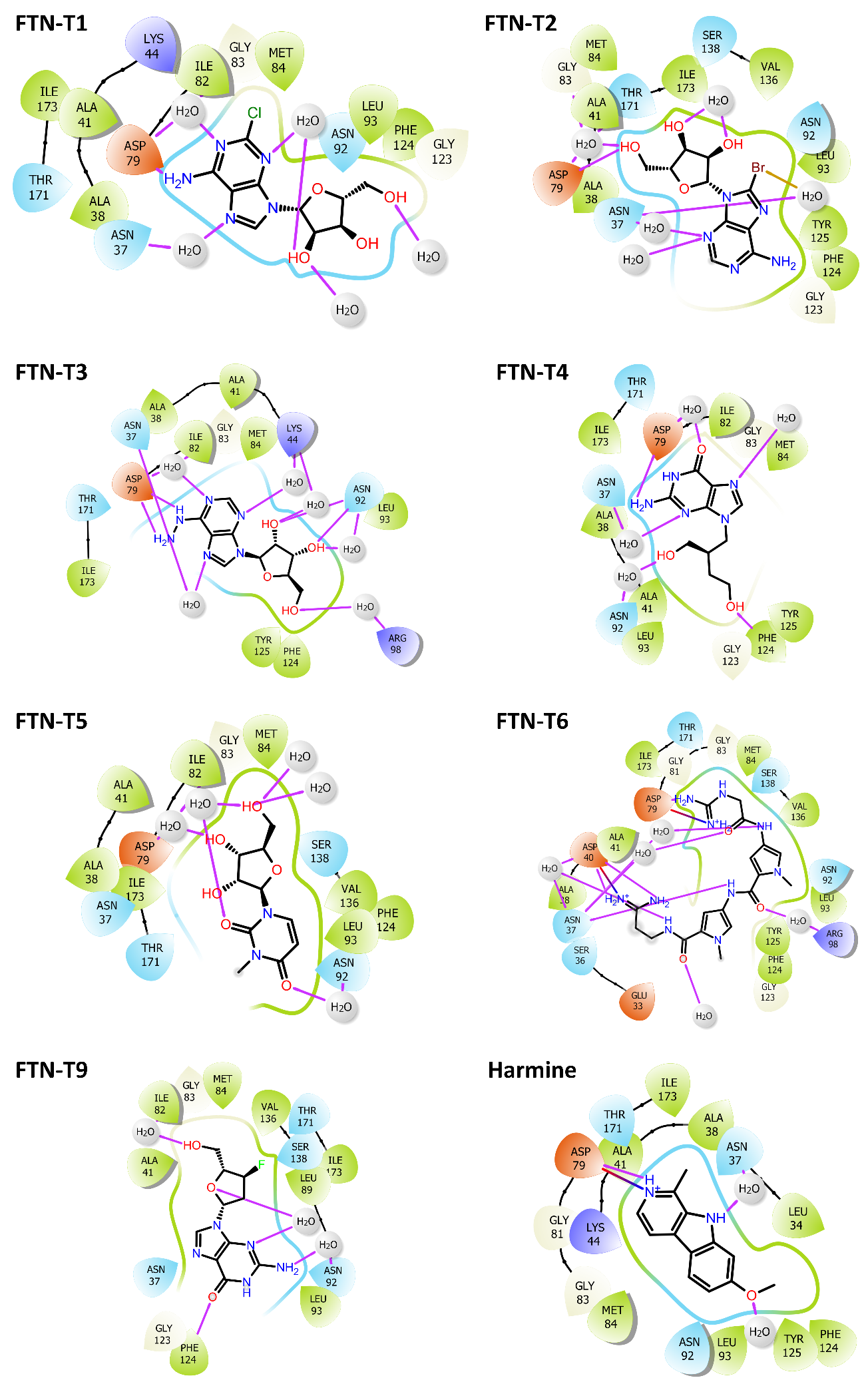

**Figure S7**:2D ligand interaction diagrams for the Induced fit docking of compounds FTN-T1, FTN-T2, FTN-T3, FTN-T4, FTN-T5, FTN-T6, FTN-T9 and Harmine against PfHsp90.

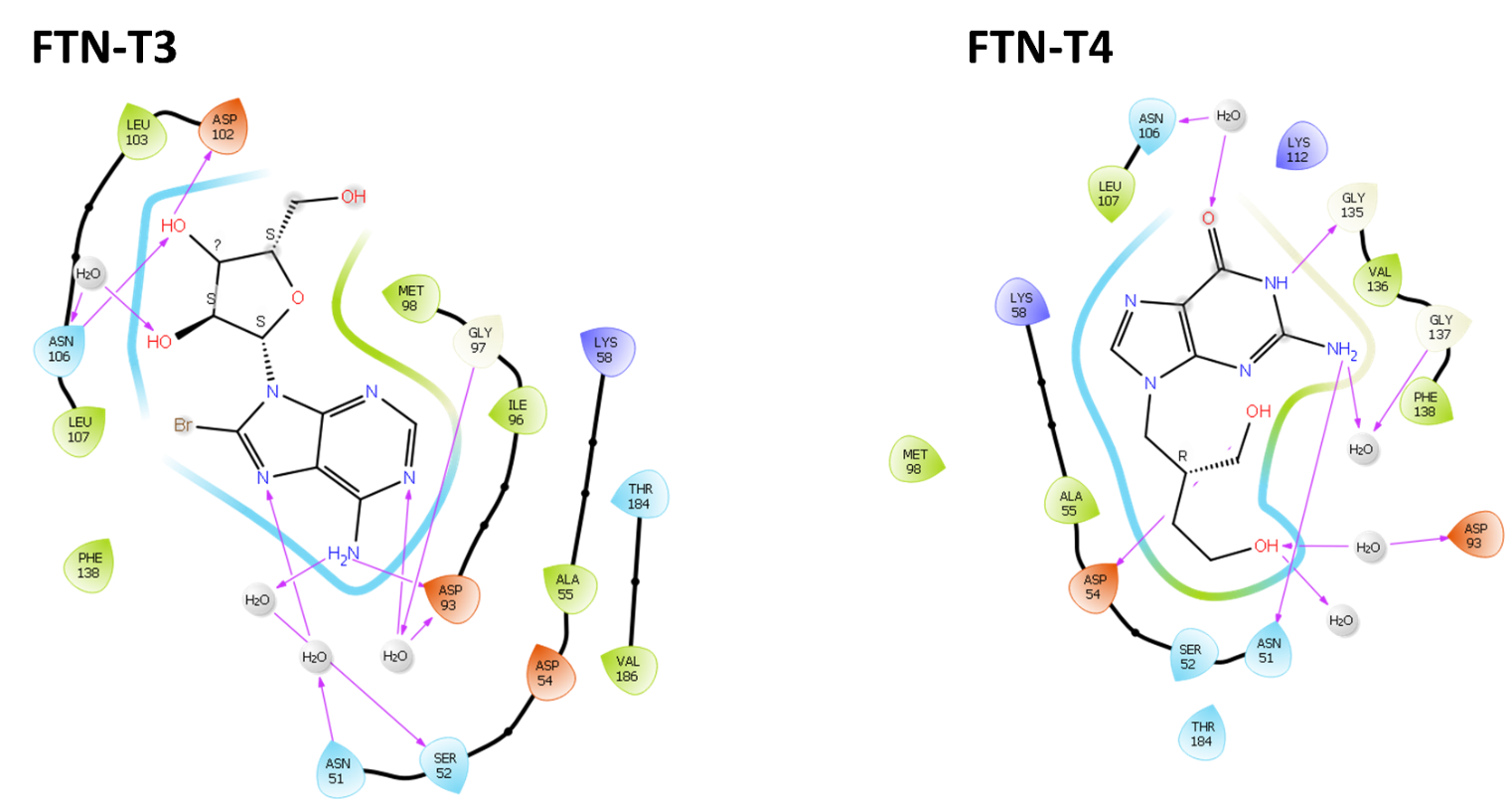

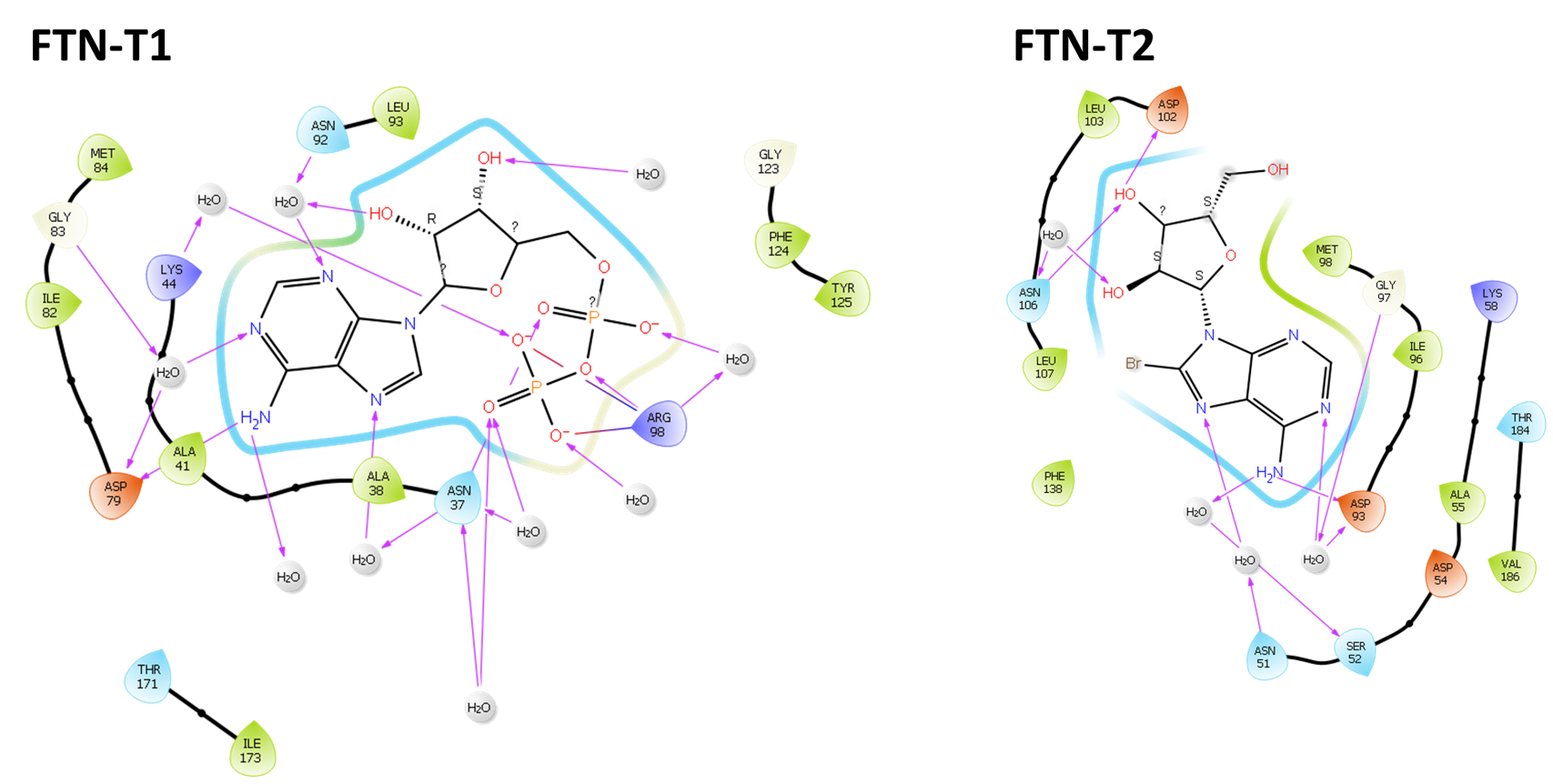

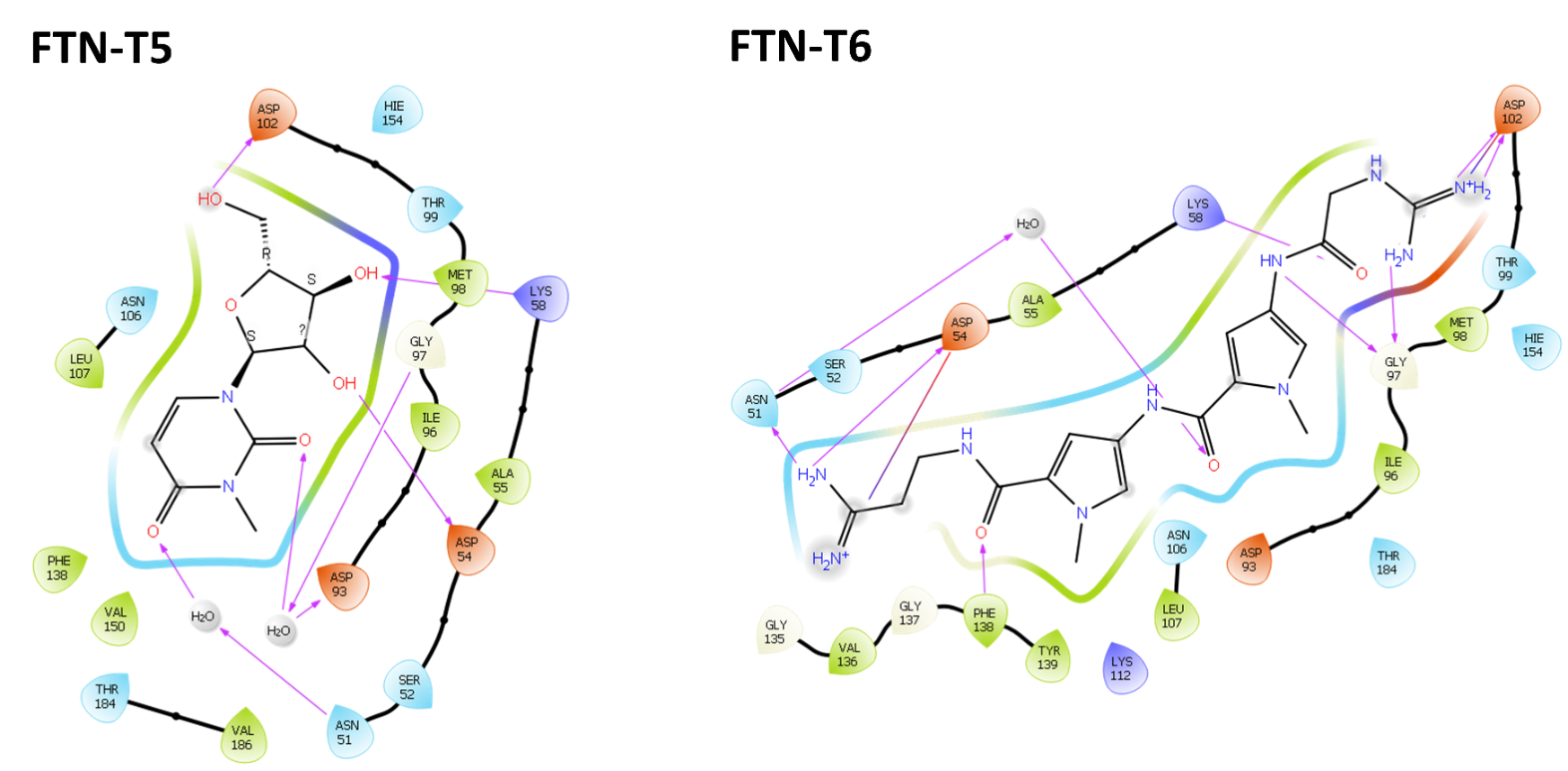

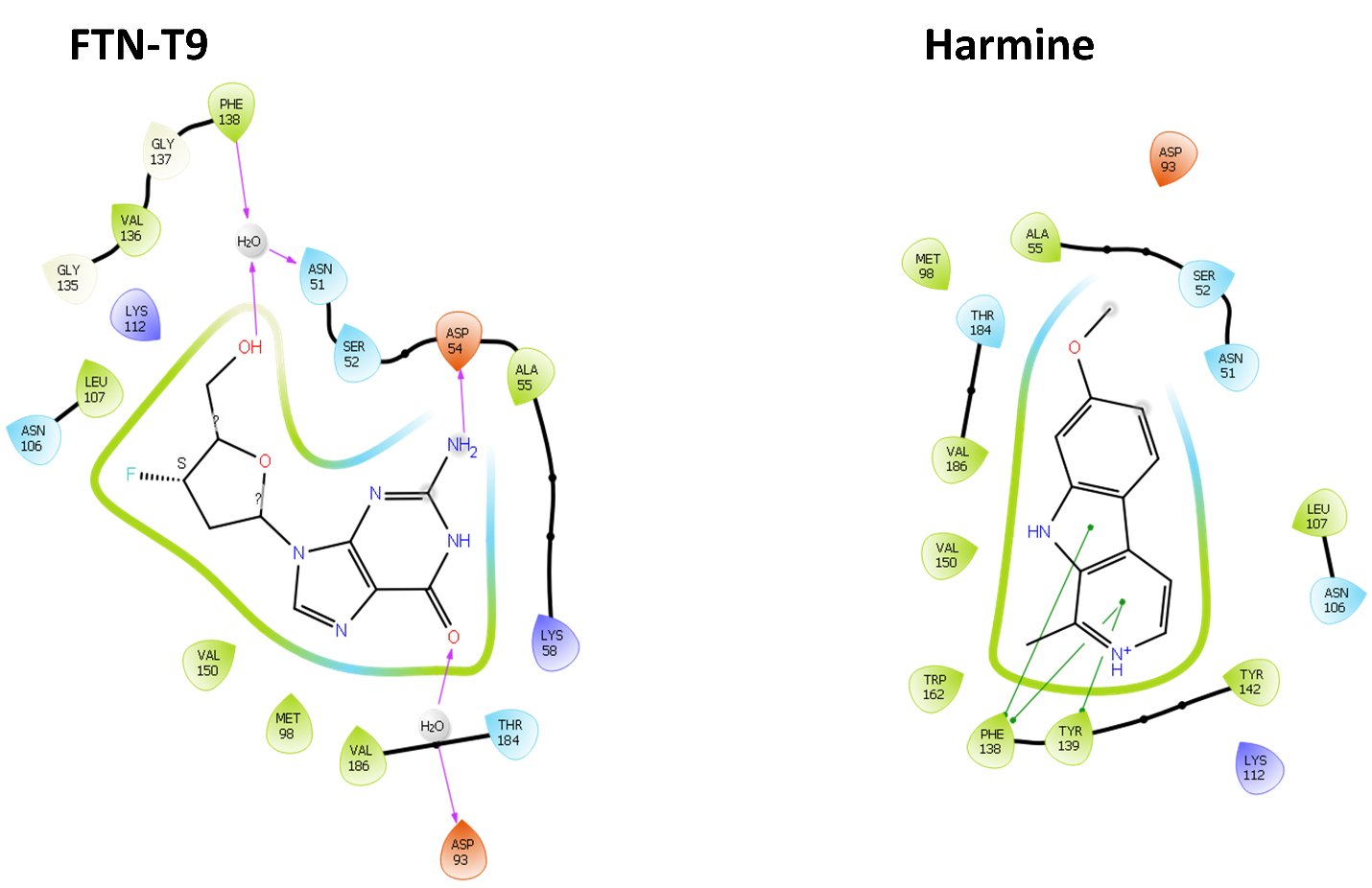

**Figure S8**: 2D ligand interaction diagrams for the Induced fit docking of compounds FTN-T1, FTN-T2, FTN-T3, FTN-T4, FTN-T5, FTN-T6, FTN-T9 and Harmine against Human Hsp90.

### D.  **ADME/T properties**

Table S5: ADME/T properties of the top 62 compounds

| Title | mol MW | QPpolrz | QPlog  Po/w | QPlogS | QPlog  HERG | QPPCaco | QPlog  BB | QPlog  Kp | HOA | PSA | Rule  Of  5 | Rule  Of  3 |
| --- | --- | --- | --- | --- | --- | --- | --- | --- | --- | --- | --- | --- |
| 40281232 | 592.696 | 66.296 | 2.890 | -7.322 | -5.272 | 12.603 | -1.499 | -4.338 | 1 | 131.590 | 1 | 3 |
| 37475529 | 593.724 | 63.129 | 5.441 | -6.796 | -5.423 | 513.824 | -1.197 | -1.431 | 1 | 115.289 | 2 | 2 |
| 315699906 | 583.692 | 66.102 | 3.077 | -6.024 | -6.025 | 44.643 | -1.646 | -4.057 | 2 | 152.695 | 2 | 2 |
| 315452139 | 593.644 | 64.297 | 3.195 | -6.684 | -5.667 | 44.406 | -2.321 | -3.953 | 1 | 188.602 | 2 | 2 |
| 48606212 | 583.643 | 60.676 | 4.420 | -6.184 | -3.901 | 19.856 | -2.182 | -2.932 | 2 | 154.053 | 2 | 3 |
| 265681303 | 552.678 | 63.066 | 3.812 | -6.170 | -6.688 | 52.291 | -1.322 | -4.514 | 1 | 134.183 | 1 | 2 |
| 314876700 | 594.712 | 65.812 | 4.556 | -6.867 | -6.839 | 132.426 | -0.989 | -3.722 | 1 | 121.105 | 1 | 2 |
| 39249872 | 517.630 | 58.734 | 2.308 | -4.557 | -5.158 | 19.826 | -1.651 | -4.984 | 2 | 144.343 | 1 | 2 |
| 41168581 | 582.701 | 64.243 | 3.291 | -6.046 | -7.054 | 31.507 | -1.946 | -4.667 | 2 | 139.510 | 1 | 2 |
| 313761635 | 552.678 | 61.555 | 2.611 | -4.131 | -7.726 | 6.099 | -1.445 | -6.775 | 1 | 142.656 | 2 | 2 |
| 4919 | 458.450 | 42.540 | -1.603 | -1.992 | -1.797 | 0.068 | -3.557 | -7.504 | 1 | 202.064 | 2 | 1 |
| 717 | 435.391 | 35.055 | -0.181 | -2.476 | -1.285 | 0.716 | -3.243 | -5.854 | 1 | 181.872 | 2 | 1 |
| 1424 | 466.496 | 49.631 | 1.349 | -4.615 | -2.969 | 0.424 | -4.517 | -7.834 | 1 | 211.092 | 2 | 1 |
| 1841 | 377.339 | 34.114 | 1.306 | -2.517 | -1.909 | 2.591 | -2.524 | -4.171 | 1 | 133.084 | 0 | 1 |
| 1424 | 466.496 | 48.971 | 1.408 | -4.553 | -2.789 | 0.734 | -4.170 | -7.490 | 1 | 200.730 | 2 | 1 |
| 4079 | 391.707 | 29.985 | -0.454 | -2.611 | -0.764 | 0.503 | -2.814 | -6.589 | 1 | 175.631 | 2 | 1 |
| 3967 | 302.229 | 22.445 | -0.923 | -1.430 | -0.549 | 0.393 | -3.121 | -6.588 | 1 | 171.012 | 1 | 1 |
| 4079 | 391.707 | 29.471 | -0.426 | -2.472 | -0.516 | 0.593 | -2.644 | -6.462 | 1 | 179.514 | 2 | 1 |
| 1133 | 332.255 | 24.474 | -1.263 | -1.194 | -0.255 | 0.521 | -3.007 | -6.381 | 1 | 176.666 | 1 | 1 |
| 1412 | 355.368 | 30.812 | -0.556 | -2.525 | -4.486 | 53.334 | -1.996 | -4.926 | 2 | 147.843 | 0 | 0 |
| 1133 | 332.255 | 25.150 | -1.186 | -1.231 | -0.265 | 0.541 | -3.002 | -6.341 | 1 | 182.509 | 1 | 1 |
| 1841 | 377.339 | 33.506 | 1.365 | -2.278 | -1.677 | 3.713 | -2.279 | -3.847 | 2 | 134.683 | 0 | 1 |
| 1715 | 346.317 | 26.817 | -2.679 | -1.858 | -4.158 | 4.580 | -2.991 | -7.006 | 1 | 189.347 | 2 | 1 |
| 3630 | 346.140 | 24.016 | -1.375 | -2.072 | -3.782 | 37.891 | -1.786 | -5.464 | 2 | 137.313 | 0 | 0 |
| 1813 | 365.328 | 33.195 | 1.263 | -2.915 | -0.905 | 2.371 | -2.193 | -5.107 | 1 | 134.786 | 0 | 1 |
| 4994 | 445.477 | 46.260 | 1.119 | -5.363 | -6.446 | 13.814 | -3.774 | -7.563 | 2 | 208.195 | 2 | 1 |
| 3885 | 371.343 | 30.645 | -1.081 | -1.971 | -0.927 | 0.279 | -3.501 | -6.761 | 1 | 195.038 | 1 | 2 |
| 4184 | 331.267 | 24.308 | -0.933 | -1.289 | -0.447 | 0.632 | -2.958 | -6.128 | 1 | 161.514 | 0 | 1 |
| 4363 | 255.233 | 20.504 | -1.925 | -1.409 | -3.741 | 16.480 | -2.453 | -5.979 | 2 | 148.929 | 0 | 1 |
| 3782 | 325.280 | 27.637 | -1.265 | -2.106 | -2.824 | 0.750 | -3.630 | -7.322 | 1 | 193.683 | 1 | 1 |
| 1261 | 368.736 | 32.641 | -0.594 | -2.752 | -4.822 | 105.397 | -1.462 | -4.462 | 2 | 138.425 | 1 | 0 |
| 3885 | 371.343 | 30.975 | -1.248 | -2.048 | -1.028 | 0.154 | -3.820 | -7.265 | 1 | 201.425 | 1 | 2 |
| 499 | 337.225 | 25.553 | -2.131 | -1.505 | -0.577 | 0.198 | -3.326 | -7.210 | 1 | 198.069 | 2 | 1 |
| 4363 | 255.233 | 20.135 | -1.952 | -1.396 | -3.844 | 17.910 | -2.450 | -5.894 | 2 | 149.810 | 0 | 1 |
| 1715 | 346.317 | 26.116 | -2.575 | -1.740 | -3.837 | 6.535 | -2.709 | -6.704 | 1 | 186.780 | 2 | 1 |
| 4843 | 273.294 | 26.363 | 0.043 | -2.603 | -4.223 | 53.635 | -1.794 | -5.134 | 2 | 117.908 | 0 | 0 |
| 499 | 337.225 | 24.571 | -2.016 | -1.413 | -0.459 | 0.352 | -3.016 | -6.694 | 1 | 197.388 | 2 | 1 |
| 3885 | 371.343 | 30.820 | -1.181 | -2.030 | -1.032 | 0.202 | -3.697 | -7.031 | 1 | 200.749 | 1 | 2 |
| 3885 | 371.343 | 31.124 | -1.282 | -2.078 | -1.084 | 0.134 | -3.906 | -7.372 | 1 | 202.789 | 1 | 2 |
| 2075 | 401.206 | 29.040 | -1.833 | -1.071 | 0.870 | 0.016 | -4.112 | -7.976 | 1 | 226.764 | 2 | 1 |
| 2925 | 364.092 | 26.700 | -3.021 | -2.143 | -2.039 | 0.172 | -2.358 | -8.496 | 1 | 157.416 | 0 | 1 |
| 3057 | 287.225 | 23.293 | -0.721 | -2.245 | -3.445 | 78.273 | -1.320 | -5.155 | 2 | 124.191 | 0 | 0 |
| 2701 | 422.232 | 27.574 | -2.017 | -0.924 | -0.256 | 0.160 | -3.242 | -7.530 | 1 | 232.169 | 1 | 1 |
| 4031 | 314.346 | 32.370 | -0.233 | -1.903 | -3.395 | 30.380 | -1.952 | -4.867 | 2 | 131.703 | 0 | 0 |
| 2474 | 297.331 | 26.155 | -0.409 | -2.314 | -3.960 | 100.185 | -1.458 | -4.619 | 3 | 115.026 | 0 | 0 |
|  | 337.245 | 26.777 | -0.639 | -3.414 | -3.127 | 0.195 | -4.470 | -8.172 | 1 | 215.135 | 1 | 1 |
| 2110 | 263.252 | 25.052 | -0.622 | -2.135 | -4.342 | 71.307 | -1.655 | -4.745 | 2 | 121.469 | 0 | 0 |
| 1589 | 235.245 | 21.188 | -0.616 | -1.798 | -3.662 | 68.549 | -1.691 | -5.076 | 2 | 119.602 | 0 | 0 |
| 2211 | 259.218 | 24.731 | 1.083 | -2.467 | -1.111 | 1.895 | -2.053 | -5.304 | 1 | 131.089 | 0 | 1 |
| 1388 | 313.206 | 26.665 | -0.970 | -1.211 | -2.057 | 37.289 | -1.111 | -4.296 | 2 | 121.524 | 0 | 0 |
| 1454 | 251.669 | 20.725 | -4.021 | 0.502 | -0.058 | 0.412 | -1.902 | -8.168 | 1 | 160.767 | 1 | 1 |
| 3550 | 268.250 | 23.523 | -0.884 | -0.793 | -4.442 | 40.922 | -0.779 | -6.292 | 2 | 109.229 | 0 | 0 |
| 3550 | 268.250 | 23.705 | -0.951 | -0.937 | -4.685 | 31.777 | -0.926 | -6.525 | 2 | 108.578 | 0 | 0 |
| 3967 | 302.229 | 22.616 | -0.919 | -1.432 | -0.514 | 0.361 | -3.135 | -6.652 | 1 | 169.562 | 1 | 1 |
| 3885 | 371.343 | 29.877 | -1.286 | -1.860 | -0.821 | 0.183 | -3.628 | -7.113 | 1 | 198.727 | 1 | 2 |
| 3550 | 268.250 | 23.734 | -0.959 | -0.995 | -4.670 | 27.745 | -0.963 | -6.666 | 2 | 113.526 | 0 | 0 |
| 1956 | 389.152 | 27.096 | -0.053 | -2.812 | -4.357 | 109.022 | -1.315 | -4.510 | 3 | 117.537 | 0 | 0 |
| 4541 | 221.218 | 20.204 | -1.124 | -1.906 | -3.947 | 42.283 | -1.928 | -5.389 | 2 | 120.798 | 0 | 0 |
| 2403 | 296.285 | 24.650 | -2.190 | -0.598 | -4.456 | 5.770 | -1.731 | -7.801 | 2 | 152.409 | 1 | 1 |
| 3816 | 298.252 | 26.263 | -1.964 | -1.564 | -4.247 | 14.786 | -2.511 | -6.154 | 2 | 167.235 | 0 | 1 |
| 95 | 272.260 | 23.326 | -1.205 | -1.551 | -4.045 | 74.627 | -1.959 | -4.751 | 2 | 134.179 | 0 | 0 |
| 845 | 251.244 | 22.294 | -1.118 | -1.739 | -3.737 | 78.966 | -1.526 | -4.860 | 2 | 119.441 | 0 | 0 |

ation on BL21(DE3) E. coli cells analysis. **(A)** SDS-PAGE stained with Coomassie blue. **(B)** Western blot analysis using His-tagged antibodies. Lane PL represents protein ladder, lane 0(J.C) uninduced cells at time point zero, lane 0 total extract of cells transformed prior to IPTG induction; lane 1 – 5 and 24 are hourly samples and overnight samples respectively, after induction with IPTG; 24(J.C) are uninduced overnight sample. (**C)** SDS-PAGE analysis of PfHsp90 protein purification samples. The purification samples were analysed by SDS-PAGE on a 12 % gel. **(D)** Western blot analysis of the purification samples. M: Molecular mass marker; Lane 2: Filtered lysate (soluble supernatant); Lane 3: Sample flow-through; Lane 4: Wash 1; Lane 5: Wash 2;lane 6: wash 3; Lane 7-9: Elution. lanes

**Figure S12**..

E. **Cross docking results**

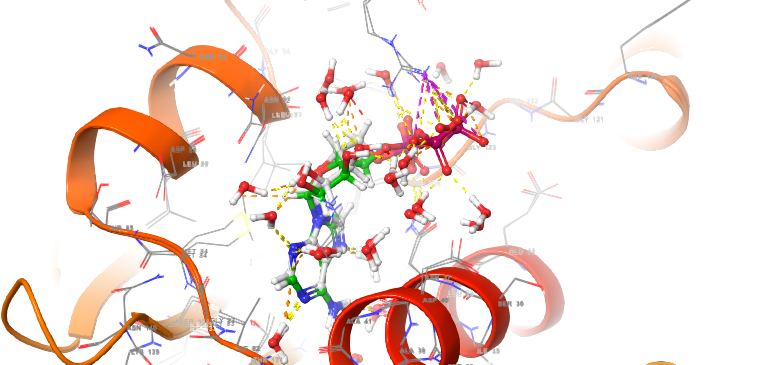

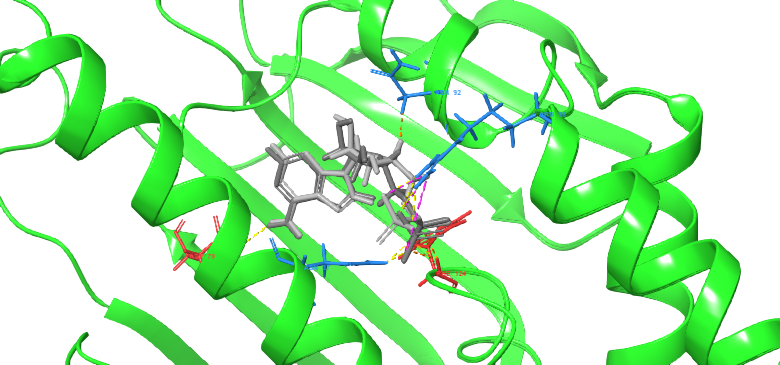

**Figure S13:** Cross docking results. To validate the docking procedure, the co-crystalyzed ligand ADP was redocked into the PfHsp90 binding site, which obtained a high degree of overlapping, as seen above with the two superimposed structures perfectly aligning with an RMSD of 0.6565 Å, indicating a reasonably accurate docking procedure.
