## Supplementary figures and images for "Auto QSAR-based Active learning docking for hit identification of potential inhibitors of *Plasmodium falciparum* Hsp90 as antimalarial agents"

### Compounds_2D_Chemical_Structures_20240424_2_TIF.tif

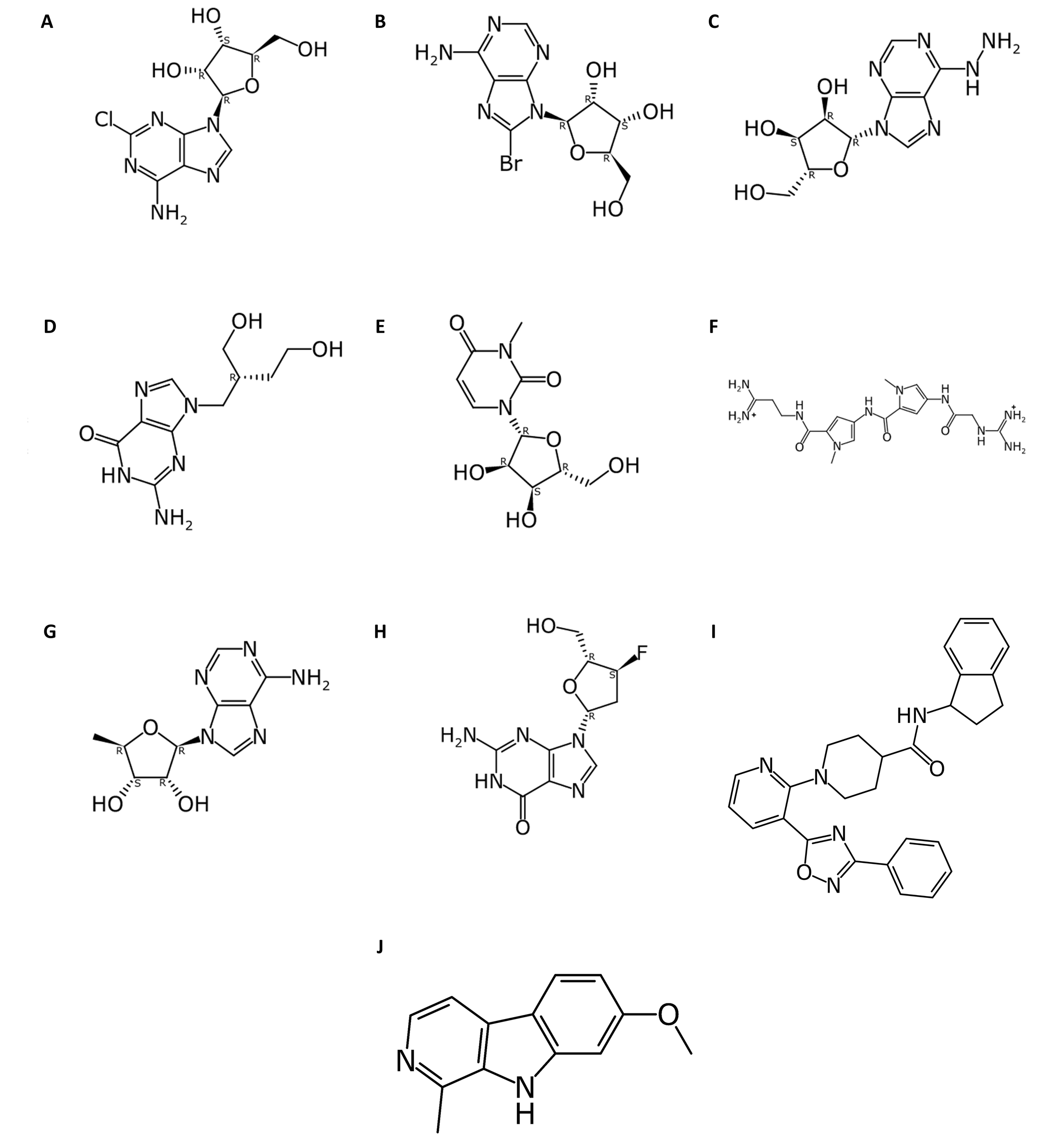
